# BOLD cofluctuation ‘events’ are predicted from static functional connectivity

**DOI:** 10.1101/2022.01.24.477543

**Authors:** Zach Ladwig, Benjamin A. Seitzman, Ally Dworetsky, Yuhua Yu, Babatunde Adeyemo, Derek M. Smith, Steven E. Petersen, Caterina Gratton

**Author notes:** Corresponding Author: Caterina Gratton.

## Abstract

Recent work identified single time points (“events”) of high regional cofluctuation in functional Magnetic Resonance Imaging (fMRI) which contain more large-scale brain network information than other, low cofluctuation time points. This suggested that events might be a discrete, temporally sparse signal which drives functional connectivity (FC) over the timeseries. However, a different, not yet explored possibility is that network information differences between time points are driven by sampling variability on a constant, static, noisy signal. Using a combination of real and simulated data, we examined the relationship between cofluctuation and network structure and asked if this relationship was unique, or if it could arise from sampling variability alone. First, we show that events are not discrete – there is a gradually increasing relationship between network structure and cofluctuation; ∼50% of samples show very strong network structure. Second, using simulations we show that this relationship is predicted from sampling variability on static FC. Finally, we show that randomly selected points can capture network structure about as well as events, largely because of their temporal spacing. Together, these results suggest that, while events exhibit particularly strong representations of static FC, there is little evidence that events are unique timepoints that drive FC structure. Instead, a parsimonious explanation for the data is that events arise from a single static, but noisy, FC structure.

**HIGHLIGHTS:**

- Past results suggested high cofluctuation BOLD “events” drive fMRI functional connectivity, FC
- Here, events were examined in both real fMRI data and a stationary null model to test this model
- In real data, >50% of BOLD timepoints show high modularity and similarity to time- averaged FC
- Stationary null models identified events with similar behavior to real data
- Events may not be a transient driver of static FC, but rather an expected outcome of it.

## INTRODUCTION

The human brain is organized in large-scale systems, or ‘networks,’ with coordinated functions such as the visual network, somatomotor network, and default mode network. In humans, these networks can be identified by grouping regions of the brain that have highly correlated spontaneous BOLD fMRI signals - regions with high “functional connectivity (FC)” (Biswal et al., 1995; Power et al., 2011; Yeo et al., 2011). These FC networks have been shown to have a canonical spatial layout (most people have the same networks represented in the same locations), with stable patterns of individual variation (each person’s network topography is slightly different from the canonical layout and consistent within themselves across time (Gordon et al., 2017; Gratton et al., 2018; Laumann et al., 2015; Seitzman et al., 2019). At both the individual and group level, functional network topology accurately predicts which regions of the brain will be activated during specific tasks (Braga et al., 2020; Gordon et al., 2017; Smith et al., 2009; Tavor et al., 2016) and variations in network topology are related to individual differences in behavior outside of the scanner (Bijsterbosch et al., 2018; Kong et al., 2019; Smith et al., 2015; van den Heuvel et al., 2009). Further, FC measured by fMRI has identified functional systems in other species which are consistent with circuit architecture measured through anatomical tracing (Du & Buckner, 2021; Margulies et al., 2009; Vincent et al., 2007).

However, analysis of spontaneous fMRI data is not straightforward. Unlike in task-fMRI, there is no predefined temporal structure that can be used to separate relevant signals from artifactual signals. Instead, typical analyses of spontaneous (resting-state) fMRI remove physiological artifacts (motion, respiration, cardiac rhythms, etc.) and assume the residual signal is the neural signal of interest (Power et al., 2020). It is typically presumed that this signal is equally present at all moments and FC is calculated using all available data over long periods.

However, recent work suggested that rather than being constantly present, FC information might be inordinately present at particular time points called “events” (Esfahlani et al., 2020). Esfahlani and colleagues found that “events,” time points with the highest BOLD signal cofluctuation, reproduce static functional connectivity patterns better than the same number of “non-events,” time points with the lowest BOLD signal cofluctuation, and require relatively few timepoints to reproduce them well. The authors concluded that rather than functional network structure being present at all timepoints, it is driven by events – a discrete and temporally sparse phenomena (Esfahlani et al., 2020).This idea has deep implications for the field: a thorough analysis of events across brain organizational levels (e.g., from systems to cellular recordings) could reveal information about the physiological mechanisms of FC and new analysis methods focused on events could improve the clinical utility of fMRI (Esfahlani et al., 2021; Greenwell et al., 2021). The idea that brain network information can be identified in a reduced data set is not new. Previous approaches such as co-activation patterns (CAPS) and point-process analysis (PPA) have been used to identify a small number of points which can capture functional network information (Liu & Duyn, 2013; Tagliazucchi et al., 2012). In this work, we consider events methods because they do not require the specification of a seed region or threshold for high-amplitude activity (Esfahlani et al., 2020).

However, there are alternative interpretations of these findings which have not yet been explored. First, it is possible that differences between “events” and “non-events” are driven by contamination in “non-events” (motion, respiration, etc.) rather than by a unique signal present during “events.” Second, it has been shown that random sampling variability in BOLD data is high and alone can create the appearance of discrete states in stationary FC simulations (Hlinka & Hadrava, 2015; Laumann et al., 2017). This principle may apply here too – sampling variability could make a subset of single points look extreme, even if they are drawn from a continuous distribution around a static FC matrix (note that if this were the case, events methodology may still be a useful way to rapidly and accurately reproduce static FC structure, but this outcome would suggest that a deep focus on events physiology relative to other timepoints has less utility). Recent work has provided a mathematical basis for how this could be the case. (Novelli & Razi, 2022). In this paper, we ask (1) if events are unique points which drive FC, (2) if non-events are unique points with high contamination, or (3) if events and non-events are an expected consequence of static FC and sampling variability.

To answer these questions, we conduct a series of analyses on real and simulated data. First, we use real data from the Midnight Scan Club dataset to test how unique events and non-events are by examining whether their properties differ markedly from intermediate timepoints. Second, we create models of simulated static BOLD data to see if sampling variability on a static signal is sufficient to explain event behavior. Finally, we examine why events are able to recreate static FC structure with so few time points.

## MATERIALS AND METHODS

### Overview and Dataset

The goal of this project was to investigate if high cofluctuation moments in resting state fMRI BOLD signals are discrete events that drive functional connectivity. We used a combination of real and simulated data for these analyses.

The publicly available Midnight Scan Club (MSC) dataset was used as our real sample dataset. The MSC dataset contains fMRI data from 10 highly sampled individuals (5 females, ages 24- 34). The data for each subject was collected across 10 fMRI sessions within 7 weeks. Across these sessions, the MSC dataset includes 5 hours of resting state fMRI; this resting-state data is the focus of our analyses. One participant (MSC08) has been excluded from these analyses because of head motion and drowsiness during rest. For single session-analysis and simulations, sessions with less than 333 usable timepoints (6/90 sessions) were excluded. All data collection was approved by the Washington University Internal Review Board and written informed consent was received from all participants. The dataset and processing have been previously described in detail (Gordon et al., 2017). A summary of relevant details is provided below.

### MRI Acquisition

MRI data were acquired on a Siemens 3T Magnetom Tim Trio with a 12-channel head coil. T1- weighted (sagittal, 224 slices, 0.8 mm isotropic resolution, TE = 3.74ms, TR = 2.4s, TI = 1.0s, flip angle = 8 degrees), T2-weighted (sagittal, 224 slices, 0.8 mm isotropic resolution, TE = 479ms, TR = 3.2s) and functional (gradient-echo EPI sequence, TE = 27ms, TR = 2.2 s, flip angle = 90, voxels =isotropic 4mm^3^, 36 axial slices) MRI images were collected. Thirty minutes of resting-state fMRI were collected at the start of each session.

### Preprocessing

Data processing for the MSC dataset is explained in detail elsewhere (Gordon et al., 2017). Relevant details for this project are shared below.

#### Structural MRI Processing

For each participant, T1 images were averaged together and used to generate a cortical surface in Freesurfer (Dale et al., 1999). These surfaces were hand-edited and registered into fs_LR_32k surface space (Glasser et al., 2013).

#### Functional MRI Processing

Slice time correction, motion correction, and intensity normalization to mode 1000 were all completed in the volume. The functional data was then registered to the T2 image (which was registered to the T1 image registered to template space), resampled to 3mm isotropic resolution and distortion corrected (Gordon et al., 2017). All alignments were applied in a single step.

#### Functional Connectivity Processing

Described in detail elsewhere (Power et al., 2014), preprocessing steps were taken to reduce the effect of artifacts on functional network analysis. This included the regression of nuisance signals (white matter, ventricles, global signal, motion and derivative and expansion terms), scrubbing of high motion frames (FD > 0.2 mm), and bandpass filtering (0.009 Hz to 0.08 Hz). For two subjects (MSC03 and MSC10), motion parameters were low pass filtered before censoring to address respiratory activity in the motion traces (Fair et al., 2020; Gordon et al., 2017). Functional data was then registered to the surface and spatially smoothed (FWHM = 6 mm, sigma = 2.55) (Marcus et al., 2011).

### Network and region definition

All analyses were done on parcellated timeseries extracted using a group-level map of 333 cortical parcels (Gordon et al., 2016). These 333 parcels can be split into 12 functional systems: somatomotor (SM), somatomotor lateral (SM-lat), visual (Vis), auditory (Aud), cingulo-opercular (CO), salience (Sal), frontoparietal (FP), dorsal attention (DAN), ventral attention (VAN), default mode (DMN), parietal memory (PMN), and retrosplenial (RSP). These systems are used to group parcels in the visualization of FC matrices.

### Comparisons between events and static functional connectivity in real data

Our first goal was to compare the network structure present in events, non-events, and intermediate bins. We followed the approach used in Esfahlani et al., 2020, calculating the RSS (root-sum-square) cofluctuation for each timepoint and binning timepoints by their RSS cofluctuation value. We compared the network structure present in each bin by creating FC matrices for each bin and calculating the similarity between bin FC and whole session FC and the modularity of bin FC. These measures are defined below.

### Cofluctuation Time Series and Events

The method for calculating cofluctuation and identifying events has been fully described elsewhere (Esfahlani et al 2020). It was followed exactly and is summarized here. The original fMRI BOLD timeseries was z-scored per parcel. For each edge (a unique pair of parcels), the z-scored values at each timepoint were multiplied, resulting in an edges X timepoints matrix. As described elsewhere, this timeseries (also called the edge-time-series), represents the exact contribution of each timepoint to static FC (Esfahlani et al., 2020). For each time point, the RSS (root-sum-square) across parcels was calculated, resulting a 1 X timepoints matrix containing the RSS cofluctuation value at each timepoint. Timepoints were binned based on RSS cofluctuation value in 5% bins, with the 5% of points with highest cofluctuation in bin one, the next 5% of points in bin two and so on.

### Functional Connectivity (FC)

For each session and subject, functional connectivity matrices were calculated using either the timepoints from the full session (‘static’ FC matrices) or from the timepoints in each bin (cofluctuation bin FC matrices). In all cases, FC was calculated by the product moment correlation between each pair of parcel timeseries, resulting in a 333X333 functional network matrix. Parcels were grouped by functional system for visualization. Edges within the diagonal blocks represent within-system correlations, and edges in the off-diagonal blocks represent between-system correlations.

### Similarity

Similarity between each bin’s FC and whole-session ‘static’ FC was calculated by vectorizing both matrices and taking the correlation between them.

### Modularity

Modularity was calculated for each bin as measure of network structure. Modularity maximization is a strategy used to arrange nodes into communities in which there are more edges within communities than expected by random chance. Each matrix was thresholded for sparseness, keeping only the top 5% of weighted edges (5% edge density). Then, all remaining edge weights were set to 1, making the graph unweighted. Newman’s spectral optimization was used to identify the optimal network structure. This structure was then quantified using Newman’s modularity statistic, Q, which measures the fraction of within-network edges minus the expected value of within-network edges in a network with the same communities but random connections (Newman & Girvan, 2004). Larger values of modularity reflect stronger community structure than expected by chance.

### Comparisons between events and static functional connectivity in simulated data

Our second goal was to test whether the relationship between network structure and cofluctuation found in real data could be explained by sampling variability in a stationary model. To examine this, we created simulated data with the same dimensionality and static covariance structure as BOLD data but sampled from a random Gaussian distribution.

### Simulated BOLD Data

For each subject and session, data was sampled from a Gaussian distribution in the dimensionality of the real data from that session. Separately, a static FC matrix was calculated from the full 30 minutes of real data. The random timeseries were projected on to the eigenvectors derived from the static FC matrix, resulting in data matched in dimensionality and covariance structure with real BOLD data but stationary by construction. This strategy is largely adapted from prior simulation work (Laumann et al., 2017). We then did the same analysis in the simulation data as was described above for real data – calculating cofluctuation, binning frames by cofluctuation, and comparing the network structure present in each bin with two measures (similarity to static FC and modularity).

### Simulated Toy Model

To aid in our second goal, we did a supplementary analysis investigating the relationship between network structure and cofluctuation in a very simple non-BOLD-like data set. The data set comprised of 4 nodes total – 2 anti-correlated networks with two nodes each. Network A was defined by the simple sine(x) wave, and both network A nodes were given that signal. Network B was defined by sine(x + π/2) and both network B nodes were given that signal. Normally distributed random noise of half the magnitude as the real signal was added to all four nodes. Then, cofluctuation was calculated for each timepoint, timepoints were binned by cofluctuation, and similarity with time-averaged FC calculated for each bin.

### PCA Analysis

To test whether simulated data would replicate the result that high amplitude cofluctuations show a particular mode of brain activity characterized by counter-fluctuations in traditionally task-positive and task-negative areas of the brain, we replicated the analysis from Esfahlani et al. 2020 in both real and simulated data. For each subject and session, we calculated a mean activity pattern for high and low cofluctuation time points (top 5% and bottom 5%). We correlated these mean activity patterns, and took the first principal component from that correlation matrix. We compared the coefficients for the high versus low points, and then mapped the coefficient scores from PC1 on to the surface of the brain.

### Temporal Spacing Analysis

Our third goal was to compare the effects of different sampling methods on the network structure present in the sampled points. We specifically wanted to investigate the effect of temporal spacing on network structure.

### Comparison of Sampling Methods

For each subject and session, we examined the network structure present in four groups of time points: high cofluctuation points (selected as the top 5% of points with highest RSS cofluctuation), low cofluctuation points (selected as the bottom 5% of points with the lowest RSS cofluctuation), consecutive points (5% of points selected consecutively beginning at a random point of the session and wrapping around when needed), and random points (5% of points selected randomly from the session). For consecutive samples, 100 iterations were done for each session to not bias the result by starting location. We further tested this by varying the number of time points selected rather than choosing 5% of time points. The number of time points was varied from 1 to 100.

### Circular Offset Analysis

In a supplemental analysis, we examined the relationship between cofluctuation and network structure after removing temporal spacing effects. To do this, we binned time points by cofluctuation and then circularly shifted them by 1-10 points in both directions to maintain the temporal spacing found in the original binning while varying their cofluctuation values. However, because we previously scrubbed high motion points from this data set, it was not possible to select 5% of time points (as in other binned analyses) and shift them without running into scrubbed points. To address this issue, we randomly sampled only 5 points per bin and used fewer bins (95-100, 85-90, 70-75, 45-50, 20-25, 0-5). This resulted in a smaller number of analyzed sessions with lower peak similarities for this analysis. To reduce bias from random sampling, we did 100 iterations and averaged the results.

### Dataset and Code Availability

MSC data has been made publicly available (https://openneuro.org/datasets/ds000224/versions/1.0.3). The parcellated timeseries used for these analyses is available here (https://github.com/GrattonLab/MSC_ROI_data).The code for the analyses in this paper is available at (https://github.com/GrattonLab/Ladwig_2022_Events_Static_FC) which will made public upon publication.

## RESULTS

### Overview

Previous work showed that moments with high amplitude cofluctuations in BOLD, or “events”, estimate static functional connectivity patterns better than low cofluctuation moments, and can do so with relatively few timepoints (Esfahlani et al., 2020). This suggested that (1) high cofluctuation events may be unique, transient phenomena which drive the large-scale network organization that we observe over long timeseries (Esfahlani et al., 2020). But there are alternate interpretations of this result: (2) differences between low and high cofluctuation could be driven by low cofluctuation timepoints exhibiting more BOLD artifacts (e.g., motion or respiration) that disrupt functional connectivity measures or (3) events may arise as a consequence of sampling from a continuous distribution, where some moments will, by chance, exhibit higher cofluctuation than others.

In this work, we test these three hypotheses. We test how network structure changes over a range of cofluctuation amplitudes, ask if this relationship is present in stationary simulated data, and analyze why events can recreate static correlation structure with so few time points.

#### 1. Network structure is continuously related to cofluctuation

First, we examined the relationship between BOLD cofluctuation and network structure across a range of cofluctuation amplitudes. Our hypotheses are visualized in **Fig. 1A.** If events are specialized discrete timepoints that drive network structure, then they should especially well represent network structure (purple) relative to other points. If low cofluctuation points are discrete timepoints more contaminated by artifacts, they should especially poorly represent network structure (yellow). If BOLD cofluctuations exhibit random variation as would be expected from sampling variation, then there should be a continuous relationship between cofluctuation amplitude and network structure (green).

**Figure 1:**
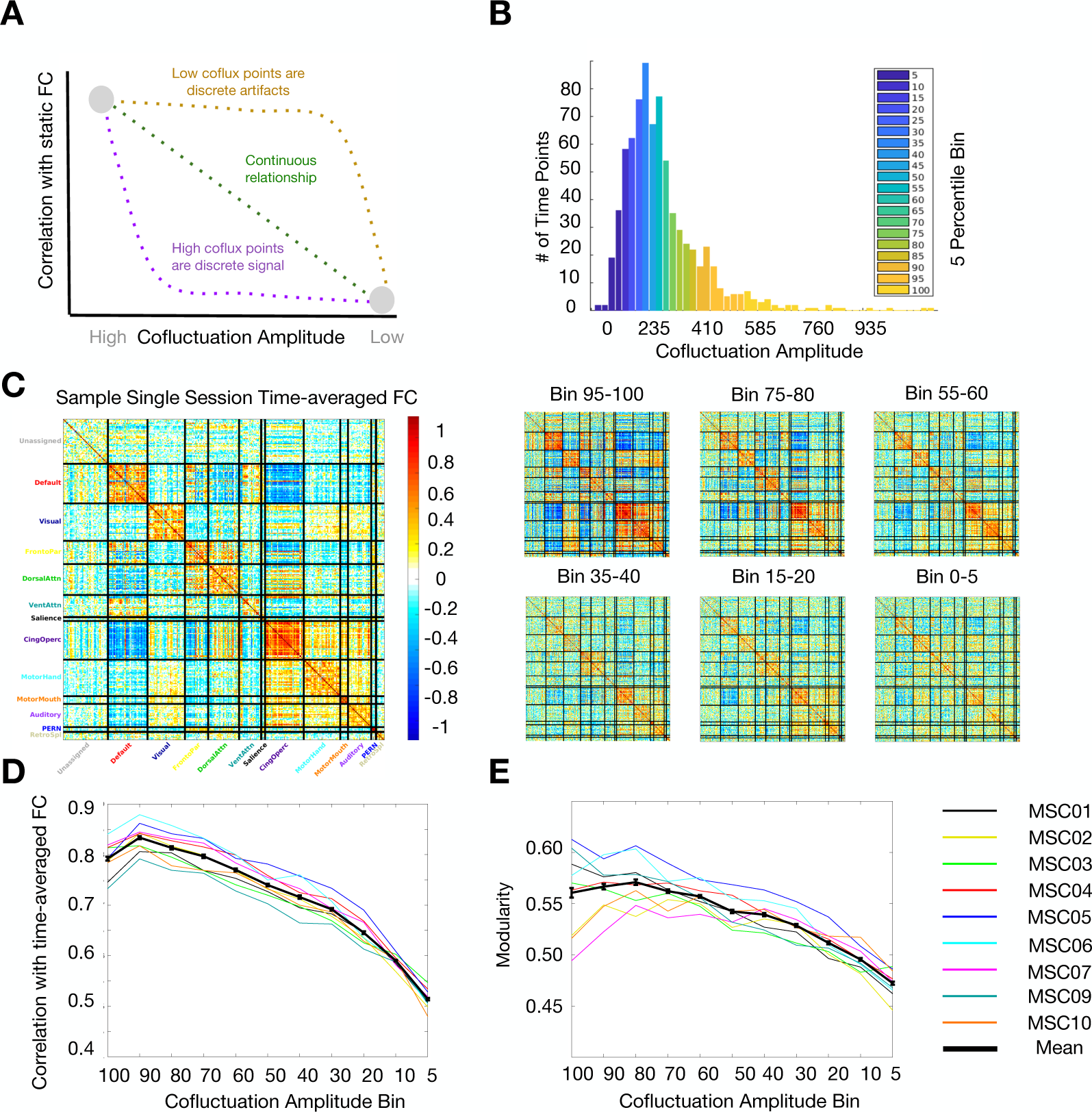
Network structure varies continuously with BOLD cofluctuation. (A) Previous literature showed that high cofluctuation events contain stronger network structure than low cofluctuation points (gray dots). We posited three hypotheses: (1) high cofluctuation points are discrete phenomena which drive network structure (purple), (2) low cofluctuation points are discrete artifacts which do not contain network structure (yellow), or (3) there is a continuous gradual relationship between cofluctuation magnitude and network structure as would be expected from sampling variability (green). (B) To test these hypotheses, we binned time points into 5 percentile bins of increasing cofluctuation. See example histogram here from MSC05 session 4. (C) For each bin, we calculated an FC matrix (examples here from MSC02 session 5) and calculated two measures of network structure – similarity to static FC and modularity. (D) Similarity to static FC increased gradually with cofluctuation for all subjects (black line = mean, colored lines = subjects, error bars represent SEM for the group). (E) Modularity increased gradually with cofluctuation as well. These results suggest that neither high nor low cofluctuation time points are discrete, unique entities.

For each participant and resting state session (30 minutes), we calculated BOLD cofluctuation amplitude at each timepoint (after standard preprocessing and denoising to improve alignment and remove artifacts, including those associated with motion, see *Methods*). In Esfahlani et al., 2020, events were defined as the top 5% of timepoints ranked by cofluctuation. We extended this, grouping timepoints in each session into discrete 5% bins based on their cofluctuation (**Fig. 1B**). For each bin, we calculated an FC matrix using Pearson’s correlation **(Fig. 1C)** and computed measures of network structure as in Esfahlani et al. 2020. (**Fig. 1D-E).**

We reproduced both results from Esfahlani et al., 2020 showing that compared to FC from the lowest cofluctuation bin (“non-events”), FC from events is more similar to whole-session FC (revents = 0.792, rlowest = 0.514, *t(89)* = 42.2, *p* = 1.2e-60) and more modular (qevents = 0.562, qlowest = 0.478, *t(89)* = 12.3 *p* = 6.0e-21) (**Fig. 1D, E).** However, when we examined the relationship across intermediate bins, we found that both metrics increased gradually with cofluctuation, not discretely for events. The increase was especially gradual at high values of cofluctuation. In fact, the top bin (events) was not substantially different than the 70^th^ percentile bin (revents = 0.792 vs. r70 = 0.797, rdiff=-0.005, *t(89)* = 0.48, p = 0.31; modularity: qevents = 0.562 vs. q70 = 0.561, qdiff=-0.001, *t*(89) = 0.15, *p* = 0.45) and only slightly different than the 50^th^ percentile bin (r50 = 0.740, q50 = 0.546, rdiff=0.052, qdiff = 0.016). Low cofluctuation points, while substantially different from events, were not obviously discrete when compared with the 10^th^ and 20^th^ percentile bins (rlowest = 0.514, qlowest = 0.478, r10 = 0.579, q10 = 0.499, r20 = 0.637, q20 = 0.515). Notably, many sets of points explicitly excluding events still recapitulated network structure well. These relationships were consistent in all 9 subjects (lines in **Fig. 1D-E**, separated by session in **Fig. S1**), suggesting that neither high nor low cofluctuation points are discrete, specialized, timepoints that drive network structure (or the lack thereof). Rather, network structure appears to be present in all bins, with variability that is positively correlated with the cofluctuation amplitude of a given time point. These results do not suggest that there are a small number of time points which drive functional connectivity. As an additional check, we re-analyzed the data omitting the low-pass filter (< 0.08 Hz) commonly applied in rsFC analysis. Low-pass filters temporally smooth data and we were concerned it may blunt the temporally discrete properties of events. We re-analyzed a single subject (MSC06) omitting the low-pass filter. As in the original analysis, it appears that network structure exists in all cofluctuation bins even in the absence of low-pass filtering (**Fig. S2**).

#### 2. Stationary simulations produce similar behavior to BOLD events and non-events

Above, we found that there was a consistent and gradual relationship between BOLD cofluctuation amplitude and network structure. Next, we asked, what drives this relationship? One possible explanation is sampling variability: with noisy data, some timepoints will have higher similarity to the session average, while others will have lower similarity, simply by chance. Here, we tested whether sampling variability could account for event behavior by creating and analyzing a simulated BOLD dataset with stationary covariance structure. In this simulated dataset, as in the real data in the previous section, we identified points of high and low cofluctuation and compared their relationship to network structure.

The procedure to generate simulated data is shown in **Fig. 2A**. For each subject and session, data was generated by sampling from a Gaussian distribution in the dimensionality of real data. This data was then projected on to the eigenvectors of the static correlation structure from the real BOLD data for that subject and session, resulting in random Gaussian data with stationary correlation structure matching real data (see *Methods*).

**Figure 2:**
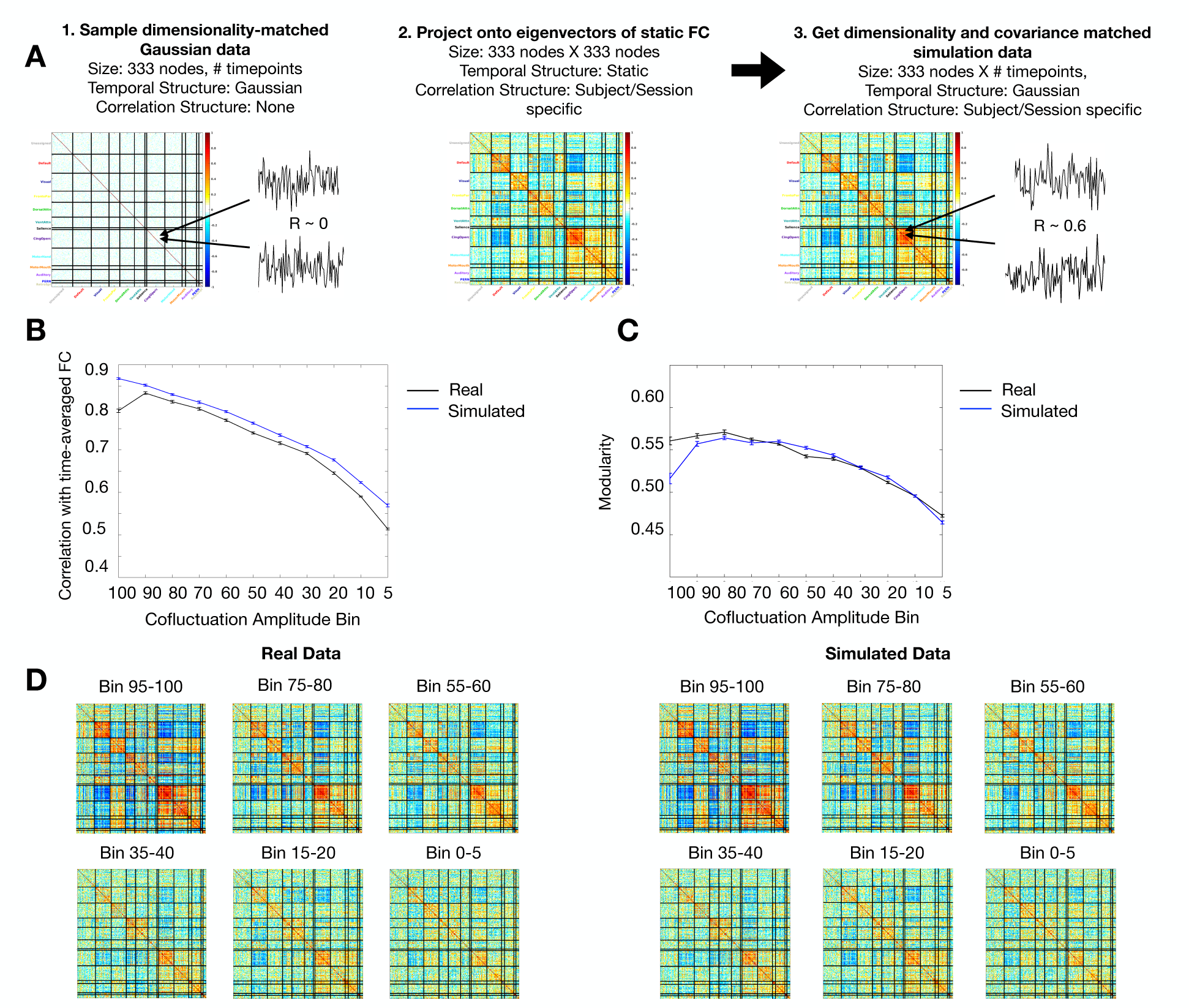
Sampling variability alone can produce event-like behavior. (A) For each subject and session, we generated a dimensionality-matched timeseries sampled from a Gaussian distribution. This time series was projected onto the eigenvectors of static FC calculated from that session. This yielded a simulated random Gaussian data set with BOLD-matched dimensionality and covariance structure. (B, C) Using the same analysis methods as in Fig. 1, we found that the relationship between network structure and cofluctuation in simulated data was remarkably similar to the one found in real data. Both (B) similarity with session FC and (C) modularity increased gradually just as they did in real data. (D) Visually, the FC matrices made from specific cofluctuation bins look similar between simulated and real data. The data shown is an example from a single session: MSC02 Session 5. These results suggest the relationship between network structure and cofluctuation amplitude can be explained by sampling variability and static FC.

The analysis from **Figure 1** was repeated on the simulated data. We calculated cofluctuation for each time point, binned time points by cofluctuation, computed FC matrices for each bin, and compared the network structure properties across bins. We found that the relationship between network structure and cofluctuation in simulated data was remarkably similar to the one found in real data. Similarity to static FC (**Fig. 2B**) and modularity (**Fig. 2C**) both showed gradually increasing relationships with cofluctuation in the simulated data, just as in real data. Visually, the network structure present in each bin was remarkably similar between simulated and real data (**Fig. 2D**). These results were consistent within individuals and sessions (**Fig. S3**). Further, we confirmed that the particular task-positive versus task-negative activity pattern associated with high amplitude cofluctuations in Esfahlani et al. 2020 was present in the simulation data as well, suggesting that this pattern, too, is sufficiently explained by static FC (**Fig. S4**). These results suggest that the difference between high and low cofluctuation moments and their relationship to network structure can be explained by sampling variability alone.

Next, we extended this simulation analysis to non-BOLD-like data by creating an extremely simple toy model made of correlated sine waves and noise **(Fig. 3A)**. Even such a basic model of correlated data shows points of high and low cofluctuation amplitude and a gradually increasing relationship between cofluctuation amplitude and network structure, similar to what is seen in BOLD and the stationary simulation. (**Fig. 3B, 3C, 3D, 3E**) This provides evidence that these properties may be the general statistical behavior of correlated timeseries and not specific to BOLD or BOLD-like data.

**Figure 3:**
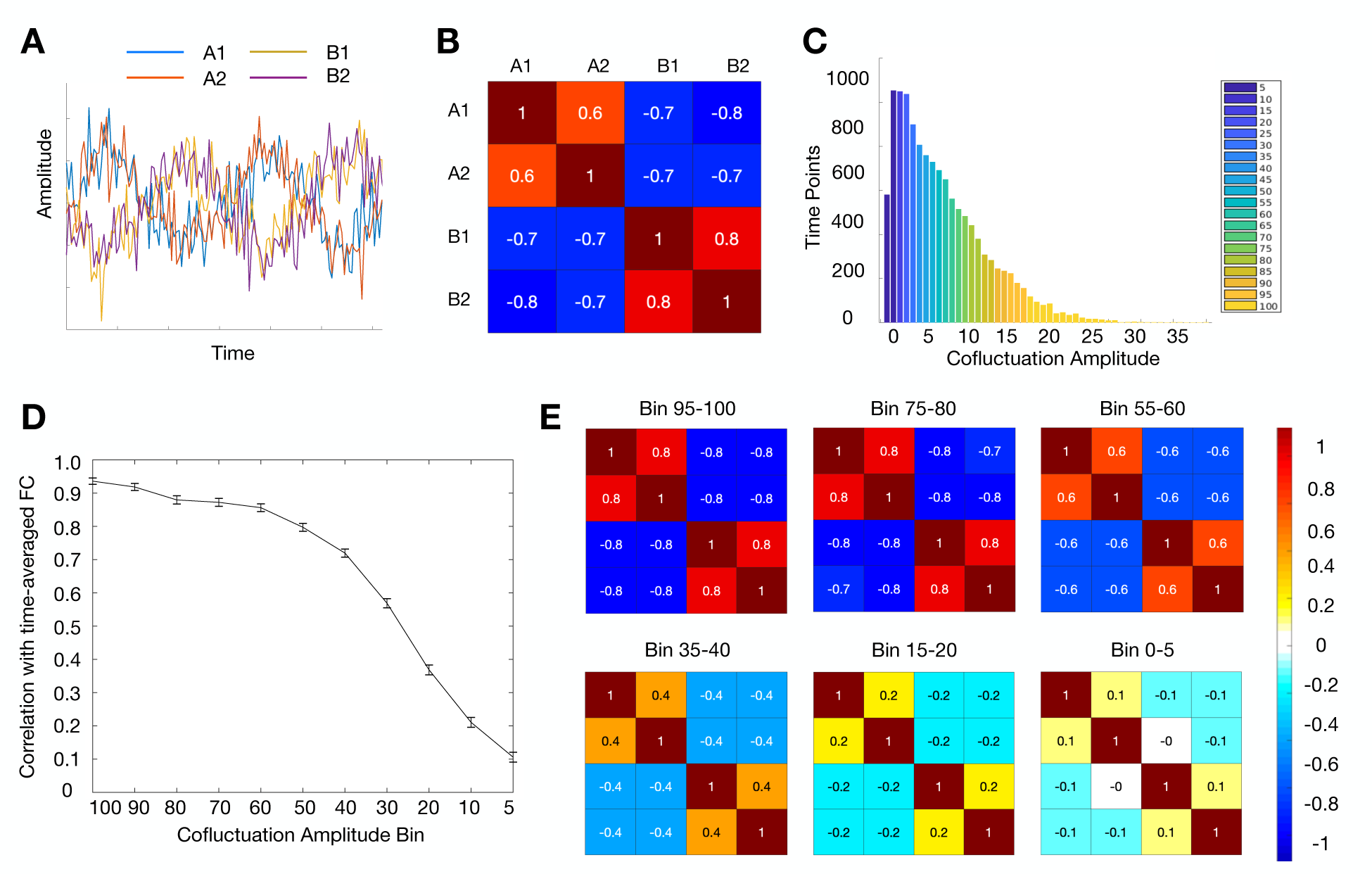
The relationship between network structure and cofluctuation is present in extremely simple non-BOLD-like models. We created a 2 network, 4 node model from sine waves and tested the relationship between network structure and cofluctuation. (A) Network A is made of two nodes, each with the sine(t) wave. Network B is made of two nodes, each with sine(t+π/2) wave. Random noise was added to all four nodes. (B) Over the time course, there is moderately high magnitude (r = 0.7) correlation between in-network nodes. (C) As in real data, there were points of high and low cofluctuation so it was possible to bin time points the same way as was done in real and BOLD-simulated data. (D) A similar relationship exists between cofluctuation and network structure where higher cofluctuation bins are better able to reproduce network structure from the overall time course. Error bars here represent SEM over 1000 iterations of the model. (E) This relationship is visually obvious in correlation matrices. In high cofluctuation bins, the two antagonistic networks are strongly present, and in low bins there is little or no relationship between nodes.

#### 3. Randomly selected timepoints can also reproduce network structure

One particularly notable property of events is their ability to recapitulate network structure with a small number of timepoints. As shown in **Fig. 1** and Esfahlani et al., 2020, 5% of time points in a 30-minute resting state session (approximately 1.5 min. of total data) show high similarity with the static FC calculated from the whole session (r = 0.792). In contrast, past work (Anderson et al., 2011; Gordon et al., 2017; Laumann et al., 2015; Noble et al., 2017) suggests that large amounts (> 30 min.) of resting state fMRI data collection are required to achieve high reliability. This discrepancy appears to bolster the suggestion that events are discrete transient phenomena which drive static functional connectivity.

However, in the previous sections, we showed that events are not unique in their ability to reproduce static FC (**Fig. 1D**). Many other points can recreate static FC. The 70^th^ percentile points were correlated with static FC at r = 0.797 and the 50^th^ percentile points are correlated at r = 0.740. These results raise the question: is the ability for a few points to recreate session FC driven by cofluctuation or something else? We hypothesized that this apparent discrepancy was related to how the events methodology samples time points. One reason that substantial data is required for reliable FC measures is because BOLD data is autocorrelated – each time point shares information with the time points around it. In contrast, the events methodology is not constrained to select temporally adjacent points. Looking at a sample timeseries, it is obvious that events are more spread out than consecutive points **(Fig. 4A).** This is confirmed by looking at the histogram of the distance between events **(Fig. 4B).**

**Figure 4:**
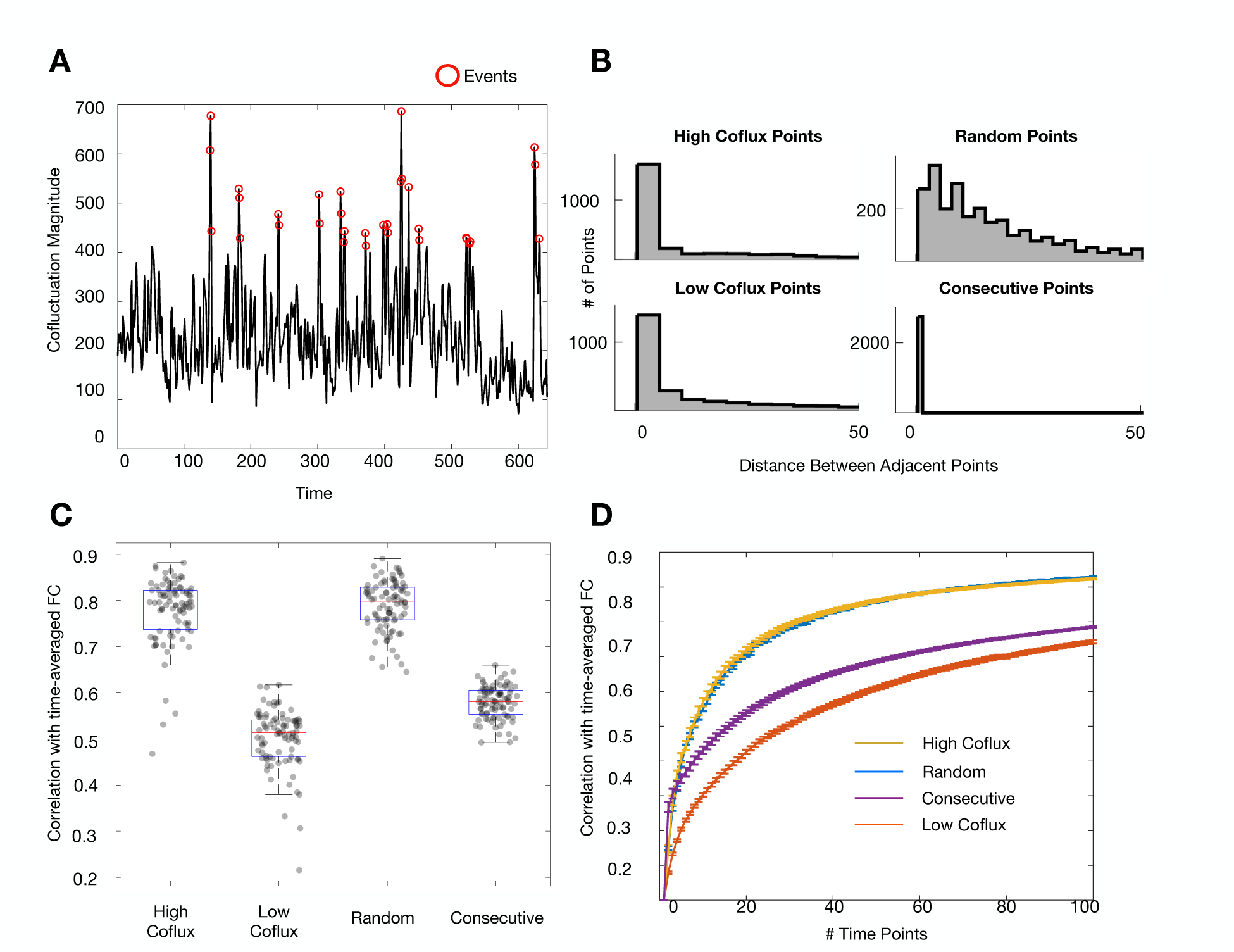
Effects of temporal spacing on estimating FC. (A) Events (red dots) are more temporally spaced than consecutive points, shown here for MSC02 session 6. (B) Histograms of distance between sampled points using consecutive, random, or cofluctuation-based sampling, aggregated over all subjects and sessions. (C) Randomly sampled points are as similar to static session FC as are events; both match static session FC much better than consecutive or low cofluctuation points. (D) These relationships hold over a range of bin sizes. These results suggest that temporal spacing is an important factor in estimating FC well.

To test the effect of temporal spacing on network structure, we compared 5% of points (a) sampled consecutively (starting from a random section of the scan), (b) sampled randomly across the whole scan, (c) sampled from the highest cofluctuation points (events), and (d) sampled from the lowest cofluctuation moments. **Fig. 4C** shows the outcome of sampling timepoints in these different ways: randomly sampled points are similarly correlated with the static session FC structure as events (r_random_ = 0.78, r_events_ = 0.79, *t(89)* = 1.7, *p* = 0.045).

Random points show substantially higher similarity to static session FC than either low cofluctuation points (r_low_ = 0.50 *t(89)* = 43.0 *p* = 1.3e^-60^), or consecutively sampled time points (r_consecutive_ =0.58, *t(89)* = 35.2 *p* = 2.42e^-54^). These results are consistent over a range of bin sizes (**Fig. 4D**), suggesting that random temporal spacing is sufficient to estimate FC well.

We note that the similarity between random sampling and event timepoints in reproducing static network FC is not immediately intuitive. One might reasonably expect that randomly selected points would fall somewhere between low and high cofluctuation points, rather than be similar to high cofluctuation points as was seen **(Fig. 4C)**. We hypothesize this unexpected result is because of temporal spacing – randomly selected points are more evenly spread out **(Fig. 4B)**, and thus better sample the underlying data, than either high or low cofluctuation points. We also examined spacing within each of the cofluctuation bins **(Fig. S5)**, finding that while all appear more spaced than consecutive points and less spaced than random samples, the extreme bins (100^th^ and 5^th^ bins) appear less spaced than the others, providing a possible explanation for why the 100^th^ bin is slightly less similar to static FC than the 90^th^ bin in real data **(Fig. 1D).** Similarity to static FC appears to be affected by two factors – cofluctuation magnitude and temporal spacing.

To disambiguate the effects of cofluctuation magnitude and temporal spacing, we circularly shifted the cofluctuation-binned time points (see *Methods*) to keep temporal spacing constant and vary cofluctuation. We found that after accounting for temporal spacing, there remained a graded hierarchy where higher cofluctuation points contained more network structure than lower cofluctuation time points (**Fig. S6).** The simulation results in the previous section suggest that this is expected and can be parsimoniously explained by sampling variability. To further investigate the impact of temporal spacing, we tested all possible spacing distances for a set number of sampled points and find that ∼10 seconds between samples is sufficient to achieve most of the benefit of temporal spacing in terms of estimating static FC **(Fig. S7).** Jointly, these findings suggest that the ability to recreate network structure with a few time points is a function of both temporal spacing (as shown here) and sampling variability (as shown in the previous section).

## DISCUSSION

In this study, we asked if “events”, time points with high BOLD cofluctuation, are discrete, transient events that drive functional connectivity. We found that events are not discrete phenomena driving FC. When they are removed, static FC structure is still present. Further, there is a gradual positive relationship between network structure and cofluctuation amplitude, with relatively similar behavior for the top 50% of timepoints, including events. Next, we asked if this gradual relationship between network structure and cofluctuation could be explained by sampling variability on static FC. We created a simulated data set matched to BOLD in dimensionality and covariance structure. Our model produced the same gradual positive relationship seen in real data, including the existence of extreme points like events, suggesting that event behavior can be explained by sampling variability alone. Finally, we analyzed why events are able to recreate static FC with so few points. We found that small numbers of randomly sampled timepoints are also able to reproduce static network structure well, suggesting that both sampling variability and temporal spacing are important factors in estimating FC. Taken together, these results support the idea that while events are an especially good representation of the network structure present in static FC, there is not evidence that they are unique points driving it.

### Should events be used to study the neural underpinnings of functional connectivity?

Although there is a large literature linking fMRI BOLD signal to neural activity (Heeger et al., 2000; Logothetis et al., 2001), the physiological mechanism of FC itself is incompletely understood. Past work suggests that BOLD FC is constrained by structural connections (Honey et al., 2009; Johnston et al., 2008; Vincent et al., 2007) and is related to correlations in neural activity (Nir et al., 2008; Shmuel & Leopold, 2008; Vincent et al., 2007) but the underlying drivers of these spontaneous activity correlations remain relatively unknown. Because events contain similar functional connectivity patterns to static functional connectivity, it was suggested that these specific moments are responsible for functional connectivity measured over the timeseries (Esfahlani et al., 2020). From a research perspective, this would make them an excellent temporal target for investigating the neural mechanism of FC.

In this work, we show that while events do match static FC well, they are not discrete markers for it. When they are discarded, static FC structure is still strongly present in the remaining time points. Further, there is a gradual and increasing relationship between cofluctuation amplitude and FC where many points (at least 50%) have a strong relationship with static FC. These results suggest that events by themselves do not (mechanistically)^1^ drive FC and it is unlikely there is a unique physiological event happening at high cofluctuation points which is creating the FC matrix. Given these observations, we consider it unlikely that investigating the unique temporal physiological activity coincident with events would glean additional new information about the physiologic origins of FC, beyond what might be seen at other timepoints as well.

However, as events show a very strong relationship to FC structure, it is possible that their study may prove useful for denoising and analysis, to provide a higher signal to noise ratio for investigations of simultaneous BOLD and direct neural recordings.

### Relationship between events and static vs. dynamic functional connectivity

The results of this work show that events can be predicted by static functional connectivity and sampling variability. However, the interpretation of these results and on the practical application of events may depend on one’s perspective about the temporal nature of FC. As has been summarized elsewhere (Lurie et al., 2020), there are two dominant perspectives on this topic. One perspective posits that functional connectivity exhibits meaningful temporal dynamics on a moment to moment basis which could represent differences in neural interactions related to ongoing cognition and task processing (Calhoun et al., 2014; R. M. Hutchison et al., 2013; Lurie et al., 2020). This view is supported by the fact that states can be found in resting-state FC data at second and minute time scales using sliding windows or instantaneous coactivation patterns (Allen et al., 2014; Chang & Glover, 2010; Petridou et al., 2013; Shakil et al., 2016), and that changes in state properties have been linked to task behavior, ongoing cognition, and arousal (Chang et al., 2016; Gonzalez-Castillo et al., 2015; M. R. Hutchison et al., 2013; Kucyi & Davis, 2014; Kupis et al., 2021; Sadaghiani et al., 2015; Tagliazucchi & Laufs, 2014) as well as more stable measures of cognitive/behavioral traits and psychiatric disease (Damaraju et al., 2014; de Lacy et al., 2017; Liégeois et al., 2019; Rashid et al., 2016). From this perspective, static FC is less significant than its constituent parts.

The second perspective posits FC is temporally stable and primarily reflects a history of co-activation between regions (Laumann & Snyder, 2021). This is supported by evidence that functional connectivity patterns are consistent within people over sessions (Finn et al., 2015; Gratton et al., 2018; Laumann et al., 2015; Miranda-Dominguez et al., 2014; Mueller et al., 2013; Poldrack et al., 2015), only slightly altered during tasks (Cole et al., 2014; Gratton et al., 2016, 2018; Krienen et al., 2014), and present in anesthesia (Mhuircheartaigh et al., 2010; Palanca et al., 2015) and slow wave sleep (Mitra et al., 2015; Sämann et al., 2011). This perspective emphasizes that resting state FC patterns are only a weak marker of ongoing cognition, and are instead more related to stable neuroanatomical constraints (Barttfeld et al., 2015; Honey et al., 2009; Lu et al., 2011), homeostatic processes (Laumann & Snyder, 2021), and learning related adaptations (Fair et al., 2007; Guerra-Carrillo et al., 2014; Lewis et al., 2009; Newbold et al., 2020; Tambini et al., 2010; Voss et al., 2012). This perspective collides with the previous one in that it suggests that the dynamic states found during rest^2^ may be explained by sampling variability, motion artifacts, and arousal (Hindriks et al., 2016; Hlinka & Hadrava, 2015; Laumann et al., 2017; Liégeois et al., 2017) rather than current cognitive content or information processing. From this second perspective, the focus of resting state analysis is on finding a clean and reliable static FC measure that may be informative about brain organization.

The information held in events, then, largely depends on which perspective one takes. From a dynamic FC states perspective, events may help identify states and characterize their properties in a more temporally specific way. Indeed, events have been used to identify states within resting state fMRI (Sporns et al., 2021), states that differentiate people (Jo et al., 2021) and states related to variation in hormone concentrations within individuals across days (Greenwell et al., 2021). However, from a static FC perspective, events may instead reflect moments of randomly good representation of the static FC structure. From this view, the previous results could be interpreted as occurring because events are particularly good timepoints for identifying stable differences between people and stable static network structure that is relevant to hormonal neurobiology.

Consistent with our findings, Novelli and Razi recently showed that many of the results of edge functional connectivity (eFC), including the presence of high amplitude cofluctuations, can be derived from static FC alone (Novelli & Razi, 2022). We showed in this current work that presence of events and the gradual relationship between cofluctuation and static FC is predictable from static FC too. While a more extended discussion of dynamic FC is outside the scope of this work, the results shown suggest that static FC and sampling variability are sufficient to explain the properties of high cofluctuation timepoints during rest reported so far. This work alone does not eliminate the possibility of multiple diverse states within resting state FC. Other modeling work has shown that events arise from biophysical models built on structural connectivity and simulated spontaneous BOLD signal dynamics (Pope et al., 2021). Indeed, it may be useful to use other simulations which include the time-varying components to see the effect of that property on events. However, the present work provides a parsimonious explanation for how events could arise from a stationary but noisy signal. We echo Novelli and Razi in our interest in future explorations of edge FC features which cannot be explained by static FC (Novelli & Razi, 2022).

Further, while we focused specifically on the events methodology in this paper, there is not an obvious reason why the results would not extend to other methods to identify single critical points like CAPS (co-activation patterns) and PPA (point-process analysis) among other similar techniques. That is, we suspect that a stationary null model of BOLD would also predict the brief instances of spontaneous brain activity reported using CAPS and PPA methodology (Liu & Duyn, 2013; Tagliazucchi et al., 2012) Indeed, recent work has shown evidence for this in both cases (Cifre et al., 2017; Matsui et al., 2021).

Finally, there is the question of the relationship between events and task-states. Previously, it was shown that events temporally synchronize across subjects during a movie watching task (Esfahlani et al., 2020). We did not explore the relationship between events and tasks in this paper, but given that arousal and tasks can create modulations in BOLD evoked signals and (more subtly) in BOLD functional connectivity patterns, we consider it possible that tasks and imposed states could change the prevalence and structure of events (Betti et al., 2013; Betzel et al., 2020; Cole et al., 2014; Gratton et al., 2016, 2018; Krienen et al., 2014; Laumann et al., 2017; Tagliazucchi & Laufs, 2014). Indeed, in the previous results it was shown that events during movies were driven by visual and attention networks rather than default and control networks shown during rest, supporting the idea that they are associated with the shared visual stimulus. However, more work is needed to fully explore these relationships and how they are separately associated with evoked signals in each region and cofluctuations.

### Practical considerations for fMRI functional connectivity analysis

Beyond fundamental neurophysiological concerns related to FC, events could be useful for a range of practical applications in FC analysis. First, we wondered if events could be used to define a filter for data points particularly suited to FC analysis. And second, given that events are good at recapitulating static FC, we wondered if it would be possible to reduce data collection by inducing more event-like time points.

Traditionally, resting state FC analyses try to isolate relevant signal by identifying and extracting known artifacts (motion, respiration, etc.) and presuming the residual data is all equally useful (Power et al., 2020). Esfahlani and colleagues’ result was particularly exciting because it suggested that, after addressing artifacts, the remaining data varied in utility for defining FC structure, with events providing a means to isolate the particularly useful components (Esfahlani et al., 2020). Although in this paper we showed that events can be explained as a consequence of sampling variability on static FC, this does not rule out that they may be a useful analytical tool. In fact, recent work has shown that ETS (edge-time-series) are better at identifying individuals than static FC (Jo et al., 2021). The strategy of seeking out points with maximal network information as a ‘denoising’ strategy is a paradigm shift in fMRI FC analysis and could be an exciting avenue of future study.

The second question is whether the fact that events can recapitulate FC with few timepoints suggests that FC may be effectively measured through much shorter data collection regimes. It has become evident in recent years that it is possible to study functional brain organization at the individual level if enough data is collected (Braga & Buckner, 2017; Gordon et al., 2017; Laumann et al., 2015; Noble et al., 2017), with most papers suggesting more than 30 minutes of high quality resting-state data is needed to measure static cortical FC reliably. This has motivated significant ongoing efforts to collect large amounts of individual ‘precision’ data (Fedorenko, 2021; Gratton & Braga, 2021; Naselaris et al., 2021; Pritschet et al., 2021) which have led to novel findings, but are costly and time-intensive, and may be difficult to acquire in clinical or pediatric populations. We wondered if, because events contain more network structure information than other time points, one could decrease data collection by increasing the rate of events and focusing analysis solely on those moments. The results in this manuscript suggest that event correspondence to static FC can be explained by sampling variability and temporal spacing - suggesting it would be difficult to ensure a high proportion of events in a short amount of data collection time. However, the results do suggest that given a sample of data, network structure could be maximized by selecting points which have high cofluctuation and are well temporally spaced (at least 10 seconds apart). Future work could explore this. In line with this, more recent publications from Betzel and colleagues have adopted an event-selection strategy which takes temporal spacing into account (Betzel et al., 2022). Beyond sampling methods, we are optimistic about new strategies for decreasing data collection needs such as new MRI techniques (Lynch et al., 2020), new parcellation strategies (Kong et al., 2019), and novel efforts to reduce artifacts (Power et al., 2020) to address these continued issues in fMRI data collection.

### Limitations

We will close by noting some limitations in this work and opportunities for future research. First. we used a dataset collected from a small number of individuals. However, we showed that the results were very similar across each participant and sessions within participants (**Fig. S1**), suggesting robustness in these results. Second, when simulating BOLD data, we used a very simple model which accounted only for spatial correlation and included no BOLD-like temporal features (e.g., autocorrelation, matched spectral structure) (Cordes et al., 2001; He et al., 2010; Liégeois et al., 2021; Zarahn et al., 1997). However, this simple model still was able to produce event-like behavior, as was an even simpler toy model from sine-waves (**Fig. 3**). That even such simple models showed event-like behavior suggests that events arise based on simple properties of the BOLD timeseries. Third, as discussed, we focused on resting-state fMRI data in this manuscript, rather than data from task sessions. However, given the synchronization of events during movie-watching, we are curious about the relationship between events and tasks, and hope to explore this in future work.

### Conclusions

In this work, we investigated high cofluctuation BOLD events and found evidence suggesting that, rather than events behaving as unique discrete timepoints that drive functional connectivity, events may arise as an expected byproduct of a static functional network structure. Event recapitulation of network structure was not unique, but varied continuously across timepoints in real data, and was present in data from which events had been excluded.

Simulations demonstrated similar responses from stationary signals. Finally, one of the primary interesting properties of events – that they can recreate static FC with a few points – is not unique and is driven in part by sampling rate. These results suggest that events are parsimoniously explained as a consequence of a highly correlated, modular, noisy signal (BOLD) and therefore might be better suited as methods for identifying good representations of static network structure than as a tool to investigate the mechanistic sources of functional connectivity.

## ACKNOWLEDGEMENTS

Funding was provided by NIH grant R01MH118370 (CG), NSF CAREER2048066 (CG), the Washington University Intellectual and Developmental Disabilities Research Center Engelhardt Family Foundation Innovation Fund (BAS) and NIH T32NS047987 (DMS). This research was supported in part through the computational resources and staff contributions provided for the Quest high performance computing facility at Northwestern University which is jointly supported by the Office of the Provost, the Office for Research, and Northwestern University Information Technology.

## CITATION DIVERSITY STATEMENT

Recent work in several fields of science has identified a bias in citation practices such that papers from women and other minority scholars are under-cited relative to the number of such papers in the field (Bertolero et al., 2020; Caplar et al., 2017; Chatterjee & Werner, 2021; Dion et al., 2018; Dworkin et al., 2020; Fulvio et al., 2021; Maliniak et al., 2013; Mitchell et al., 2013; Wang et al., 2021). Here we sought to proactively consider choosing references that reflect the diversity of the field in thought, form of contribution, gender, race, ethnicity, and other factors. First, we obtained the predicted gender of the first and last author of each reference by using databases that store the probability of a first name being carried by a woman (Dworkin et al., 2020; Zhou et al., 2020). By this measure (and excluding self-citations to the first and last authors of our current paper), our references contain 8.03% woman(first)/woman(last), 10.87% man/woman, 19.14% woman/man, and 61.96% man/man. This method is limited in that a) names, pronouns, and social media profiles used to construct the databases may not, in every case, be indicative of gender identity and b) it cannot account for intersex, non-binary, or transgender people. Second, we obtained predicted racial/ethnic category of the first and last author of each reference by databases that store the probability of a first and last name being carried by an author of color (Ambekar et al., 2009; Sood & Laohaprapanon, 2018). By this measure (and excluding self-citations), our references contain 11.74% author of color (first)/author of color(last), 13.99% white author/author of color, 27.03% author of color/white author, and 47.24% white author/white author. This method is limited in that a) names and Florida Voter Data to make the predictions may not be indicative of racial/ethnic identity, and b) it cannot account for Indigenous and mixed-race authors, or those who may face differential biases due to the ambiguous racialization or ethnicization of their names. We look forward to future work that could help us to better understand how to support equitable practices in science.

## SUPPLEMENTAL INFORMATION

**Figure S1, related to Figure 1.**
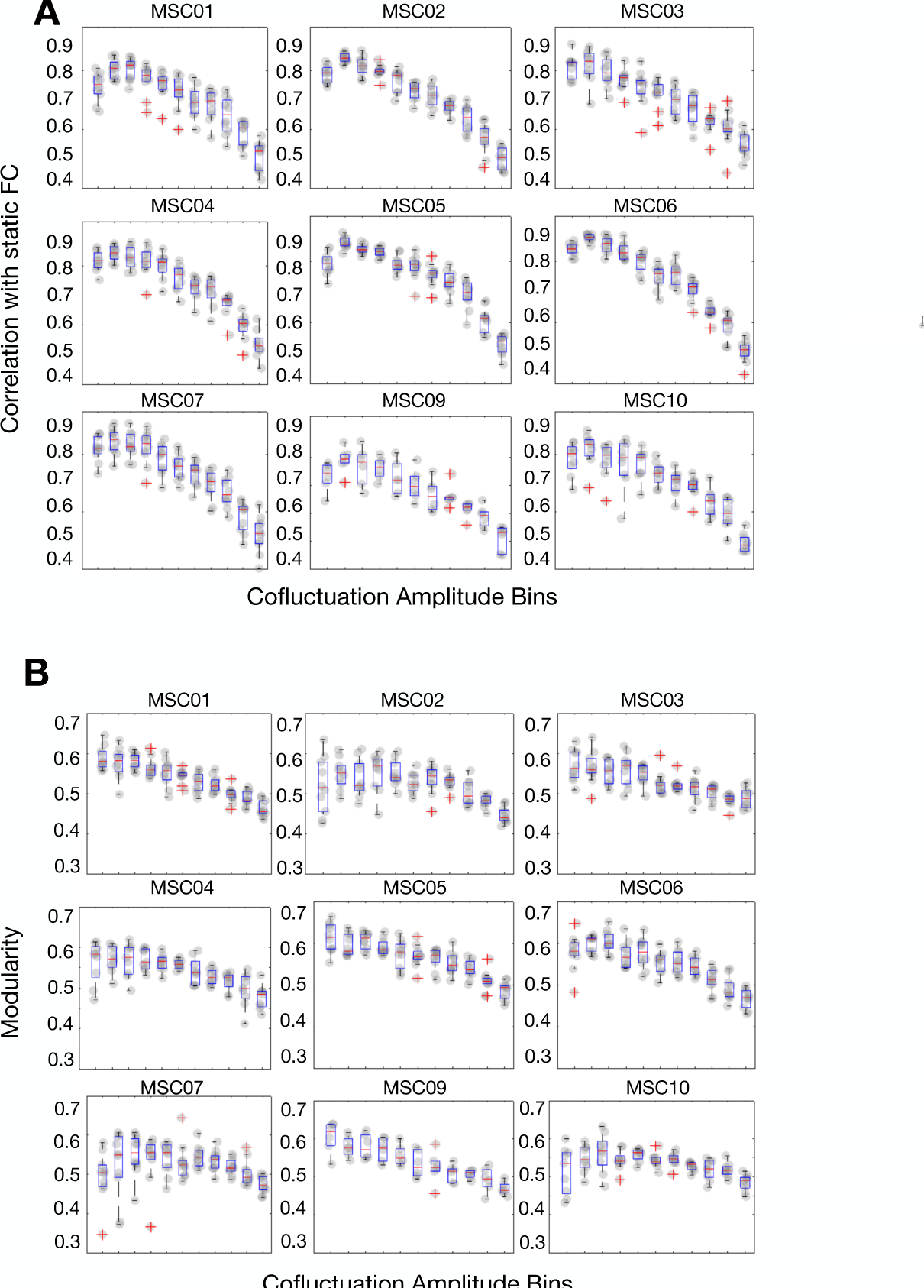
Individual subject results for Figure 1 analyses. In all subjects and sessions, the relationships present between cofluctuation and correlation with time-averaged FC (A) and modularity (B) are continuous, positive, and gradual. It suggests that in all cases, neither high or low cofluctuation time points are discrete. Boxplots show the median, 25^th^ and 75^th^ percentile values per subject calculated across sessions.

**Figure S2, related to Figure 1.**
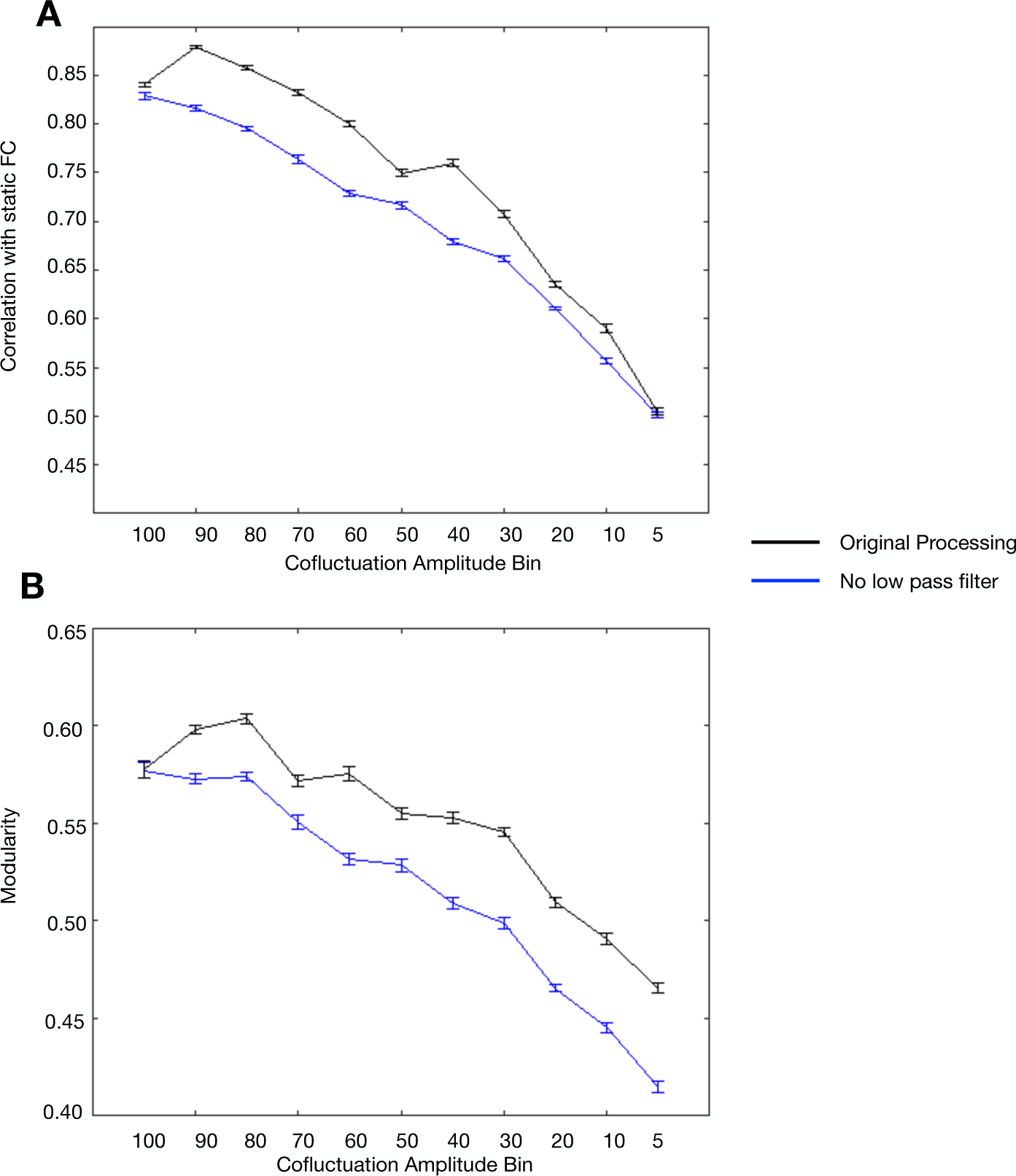
Re-analysis of MSC06 data without a lowpass filter. The general pattern of increasing network structure with cofluctuation is still present in non-low-pass-filtered data across 10 sessions in one subject. This is reflected in correlation with static FC (A) and modularity (B). Notably, there is an overall decrease in network structure across all bins versus filtered data. This is to be expected, as high frequencies of BOLD contain noise (e.g., respiratory and cardiac) that may obscure network structure.

**Figure S3, related to Figure 2.**
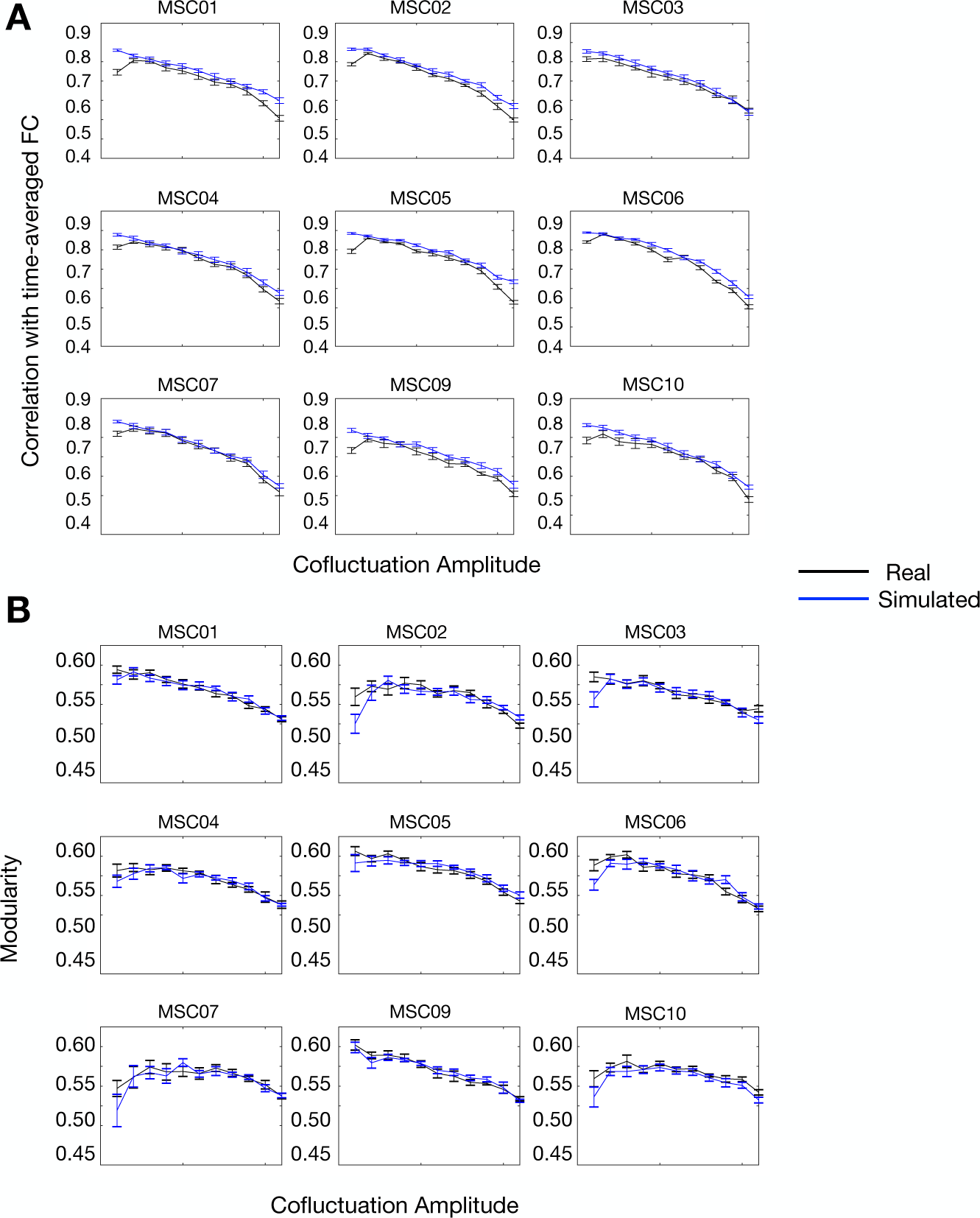
Individual subject results for simulation analysis. In all subjects, the relationships present between cofluctuation amplitude and correlation with time-averaged FC (A) and modularity (B) are continuous, positive, and extremely similar to the relationships present in real data. This suggests sampling variability in static FC is sufficient to explain the presence of high and low cofluctuation points in real data.

**Figure S4, related to Figure 2.**
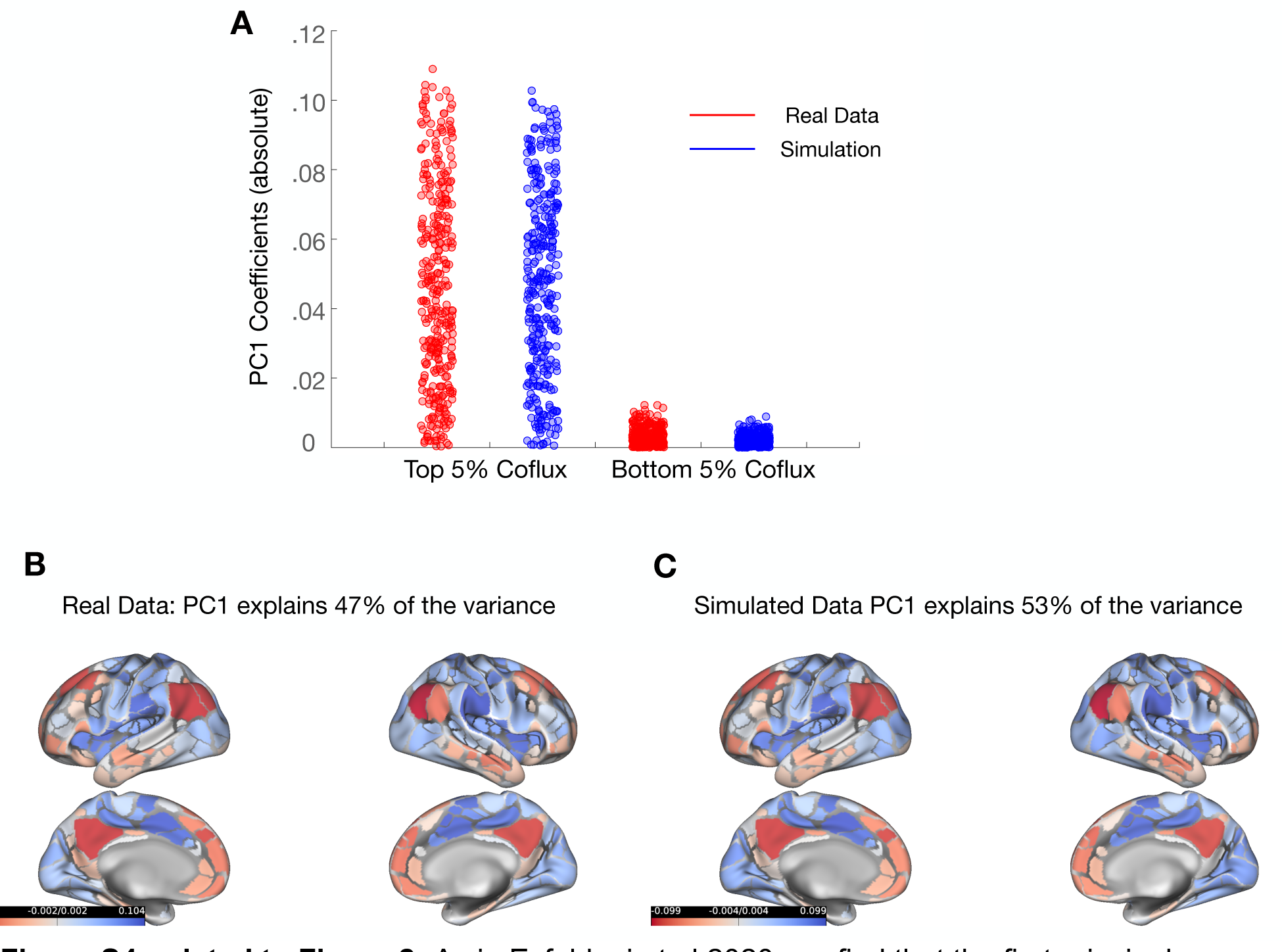
As in Esfahlani et al 2020, we find that the first principal component extracted from high and low amplitude cofluctuations corresponded to an activity pattern with shared fluctuations between default-mode and control areas and anti-correlated with fluctuations in sensorimotor and attention areas (A). We find these patterns are nearly identical in simulated data (B), suggesting that static FC is sufficient to explain this result.

**Figure S5, related to Figure 4.**
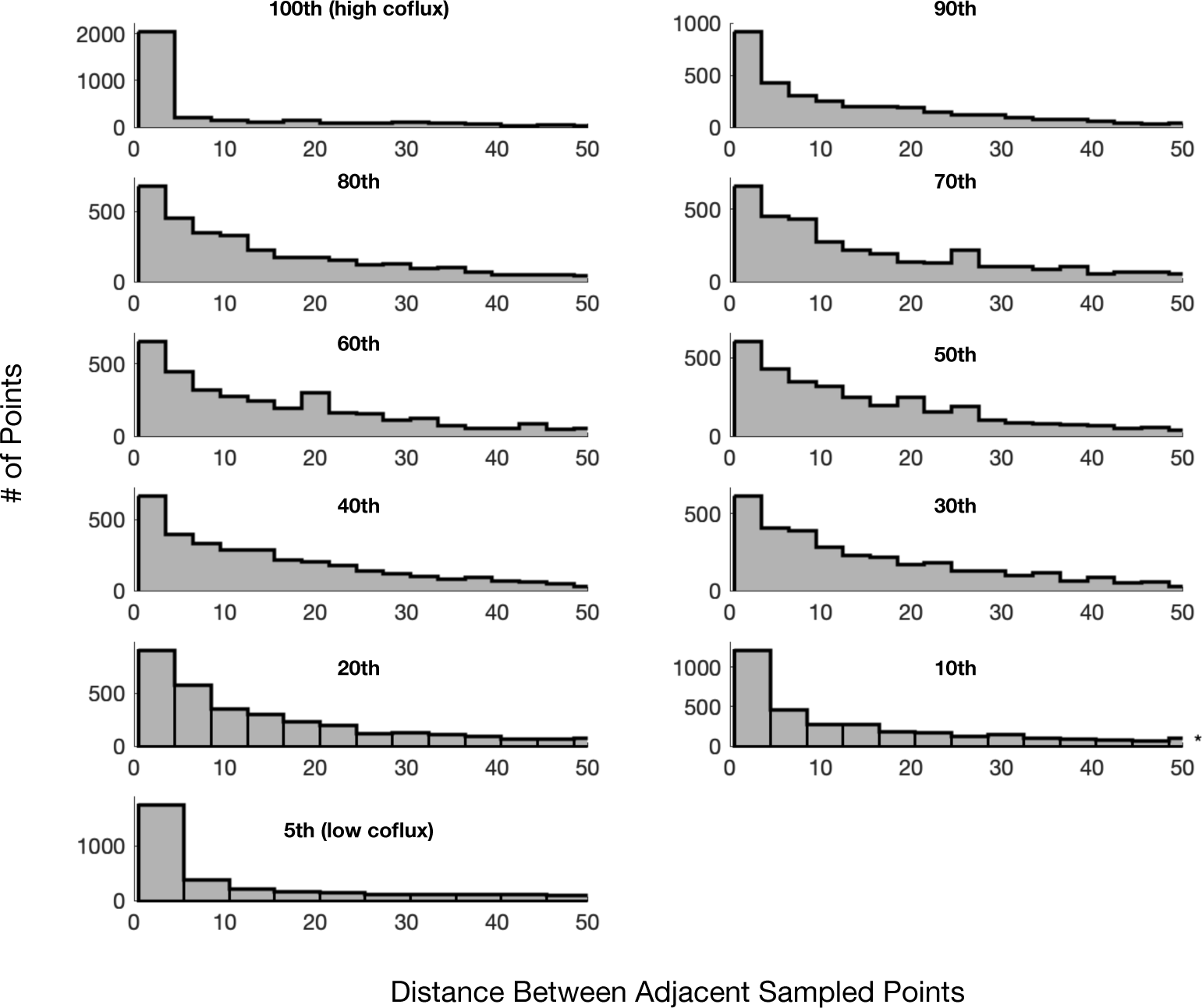
Temporal spacing histograms as in Fig. 4B but for 11 cofluctuation bins. Temporal spacing for all bins appears more spaced than consecutive and less spaced than random. It appears less spaced for the most extreme (100^th^ and 5^th^) bins than for other bins.

**Figure S6, related to Figure 4.**
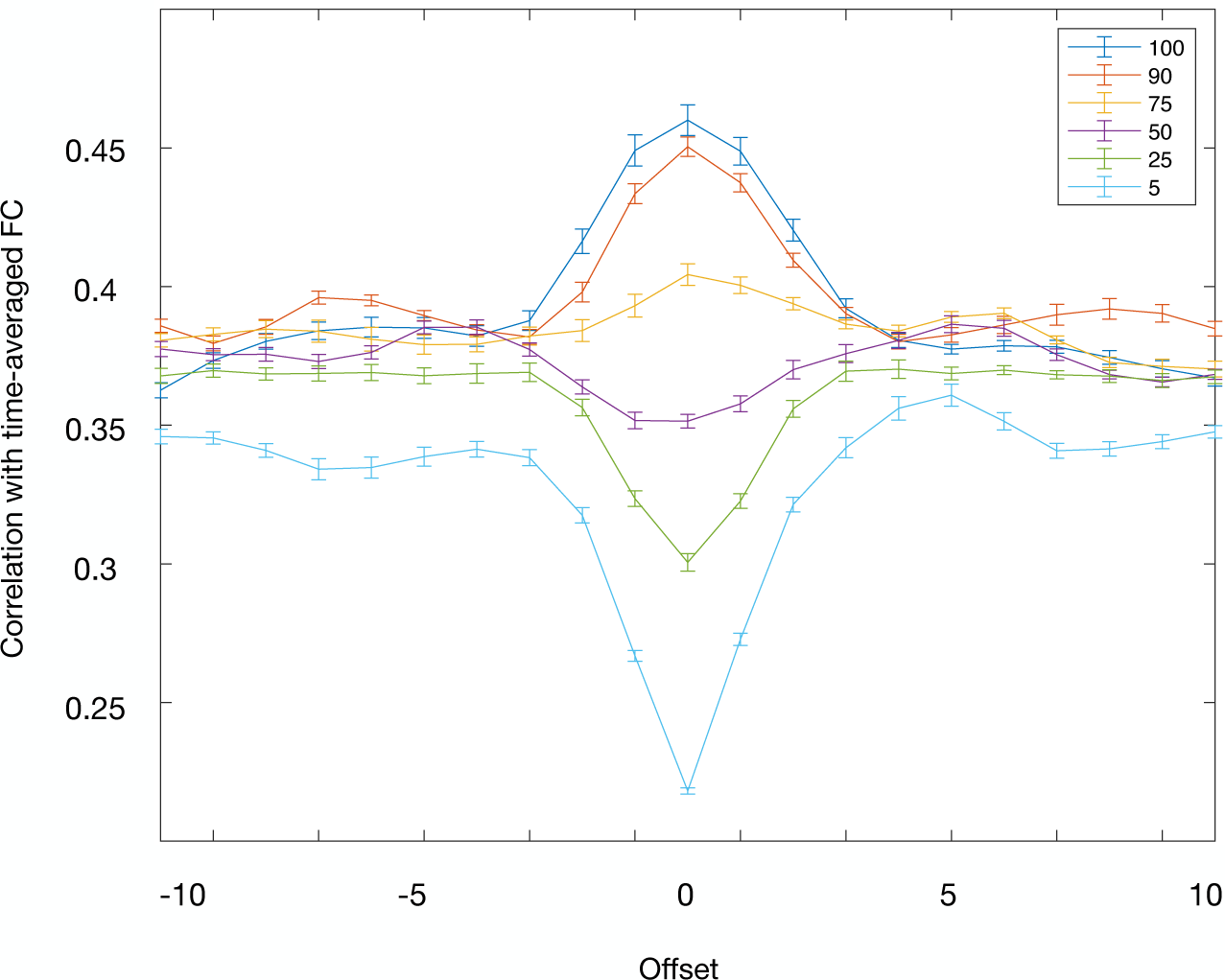
After accounting for spacing, there remains a graded hierarchy of network structure with cofluctuation. We completed a circular shifting analysis (see *Methods*) where time points are binned by cofluctuation and then circularly offset. Due to scrubbing, it was not possible to select as many time points per bin (typically all time points, 5% of total time points) without running into scrubbed points while circularly shifting the values. To address this issue, we randomly sampled only 5 eligible points per bin and used fewer bins (95-100, 85-90, 70-75, 45-50, 20-25, 0-5). This resulted in a smaller number of sessions (53/90) which contained full data for all bin and shift combinations. Because we used only five points, correlation values (maximum = 0.45 in group data) were much lower than when we calculate similarity with session FC using 5% of available points. We completed 100 iterations of this analysis and averaged the results to reduce single trial bias of the random selection of 5 eligible points. Higher cofluctuation bins have stronger correlation with time averaged FC compared to their offset counterparts and lower cofluctuation bins have weaker correlation. This suggests that while temporal spacing does in part drive the similarity of events to static network structure, there is a relationship between cofluctuation and network structure. Fig. 2 suggests this is parsimoniously explained by sampling variability.

**Figure S7, related to Figure 4.**
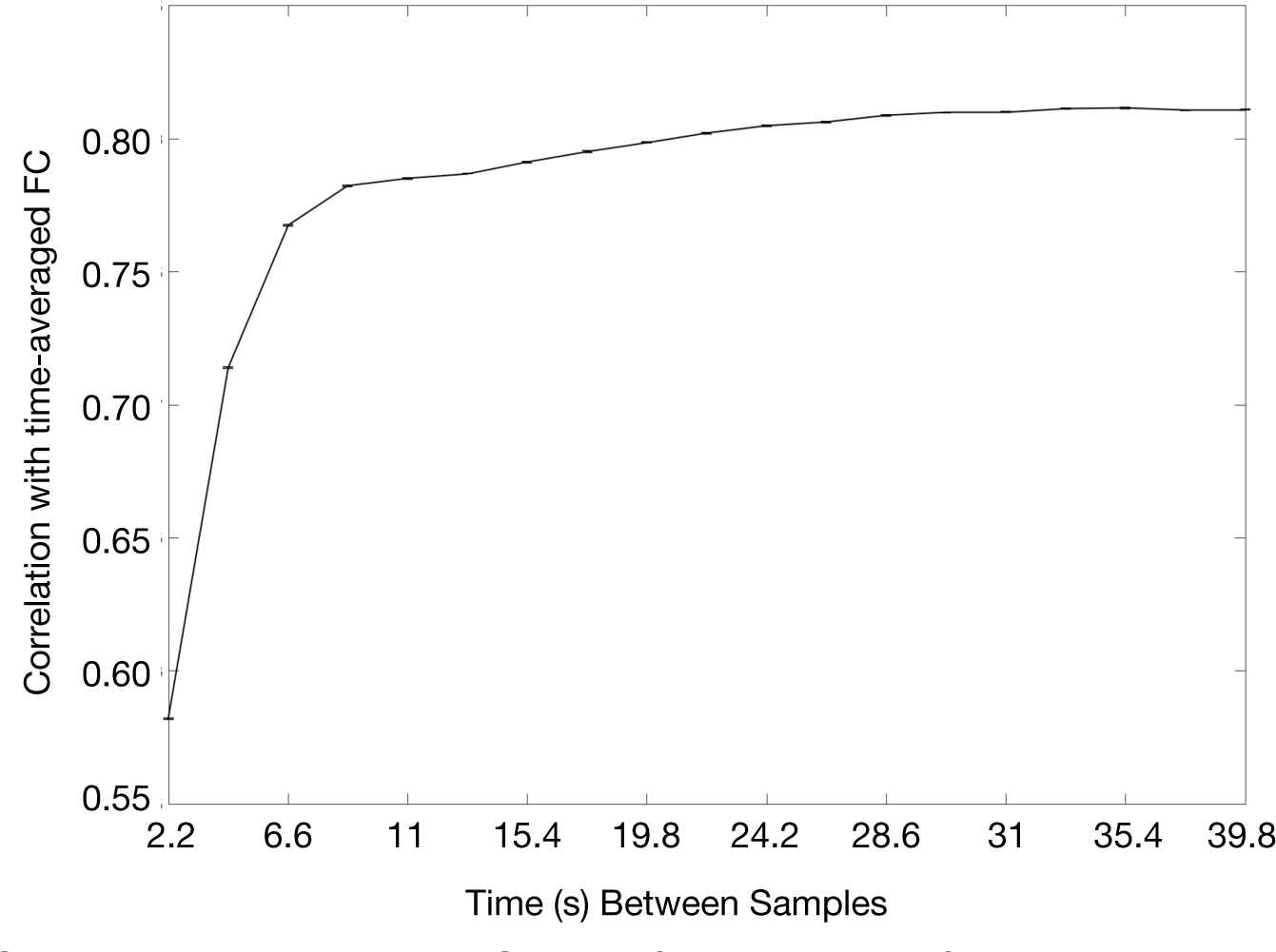
Given a fixed number of sampled points, increasing temporal spacing between samples increases the similarity with static FC. We sampled 5% of time points for each session in each subject and correlated it with static FC. We iterated over sampling spacing from sampling consecutive points (0 points between samples) to the maximum possible spacing (total # time points / # sampled points). We find that ∼10 seconds (4-5 TRs here) between samples appears adequate to achieve most of the benefit of spaced sampling.

## Footnotes

1 Events do not appear to drive FC in a unique way but do contribute the most to FC estimates as a mathematical necessity of their definition and relationship with correlation.

2 Significant differences in dynamic FC states are seen during tasks, but they tend to be relatively small (Cole et al., 2014; Gratton et al., 2018; Krienen et al., 2014; Laumann et al., 2017).

